# Early Aβ-induced changes in the enteric nervous system and the gut: structure, function, and motility

**DOI:** 10.1101/2025.09.26.678722

**Authors:** Manuela Gries, Hannah Puhl, Steven Schulte, Anne Christmann, Stephanie Rommel, Marko Baller, Monika Martin, Tobias Hartmann, Marcus O.W. Grimm, Karl-Herbert Schäfer

## Abstract

Alzheimer’s disease is increasingly recognized as affecting not only the central nervous system but also the autonomic nervous system, comprising the enteric nervous system, and thus the gut. Given the structural and functional similarities between the enteric and central nervous system, including susceptibility to Aβ, this study investigated the early and acute effects of monomeric Aβ on primary enteric neurons *in vitro* and on intestinal motility *ex vivo*. Acute Aβ application caused marked neuronal hyperexcitability, with increased spike frequencies and calcium influx, and led to enhanced intestinal contractility without altering frequency or timing. Prolonged 72-hour exposure did not induce cell death or apoptosis but significantly reduced neurite outgrowth and decreased synaptic markers and beta II tubulin. These findings suggest that acute Aβ primarily drives the excitability of enteric neurons and intestinal motility, while prolonged exposure subsequently leads to structural and synaptic changes. Overall, the study points to early dysfunction of the enteric nervous system in Alzheimer’s disease and highlights the gut as a potential target for early interventions.

**Synopsis:** 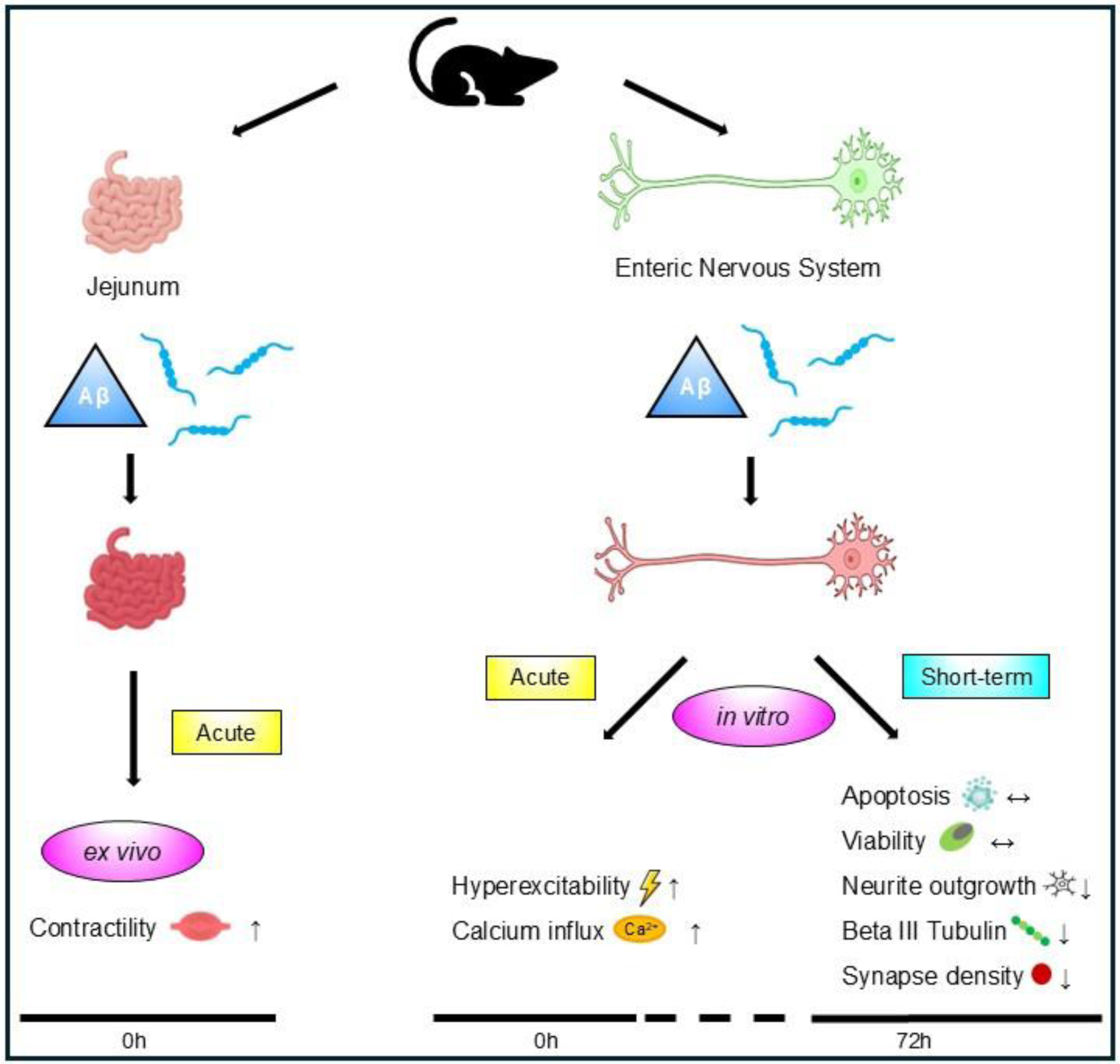

The pathology of Alzheimer’s disease is increasingly recognized in the enteric nervous system (ENS), suggesting an early contribution of the gut to the onset of the disease. This study investigated the acute and short-term effects of amyloid-β (Aβ) on intestinal motility and enteric neurons to elucidate early functional and structural changes beyond the brain.

- Acute Aβ exposure enhanced intestinal contractility ex vivo.
- Enteric neurons showed hyperexcitability and increased Ca²⁺ influx.
- After 72 h, neuronal viability and apoptosis remained unaffected.
- Prolonged exposure reduced neurite outgrowth and βIII-tubulin expression.
- Synaptic density was significantly decreased.

## Introduction

Alzheimer’s disease (AD) is a progressive, age-related neurodegenerative disorder and the most common cause of dementia, affecting more than 50 million people worldwide (Yu *et al*, 2025). Brain pathology is characterized, as one hallmark, by the accumulation of neurotoxic extracellular beta-amyloid (Aβ) peptides, which play a key role in the progression of the disease (Hardy & Higgins, 1992). These peptides are associated with disrupted neuronal activity and network dysfunction, ultimately leading to neuronal death (Palop, 2011a). Beyond Aβ plaques, impaired neurogenesis, reduced synaptic plasticity, and widespread synapse loss are prominent features (Palop, 2011b; Selkoe, 2002). Small, soluble Aβ oligomers are considered the main drivers of toxicity (Walsh & Selkoe, 2007), whereas monomers appear nontoxic or even neuroprotective at low concentrations (Giuffrida *et al*, 2009). Under pathological conditions, monomers aggregate into toxic oligomers, then into protofibrils and insoluble fibrils, forming the core of amyloid plaques (Penke *et al*, 2020a).

Synaptic degeneration likely begins early in AD, supported by evidence from patients with mild cognitive impairment (Masliah *et al*, 2001; Scheff *et al*, 2006). This deterioration may start as reversible functional decline and progress to irreversible structural loss (Rapoport, 1999). Even neurons that survive may lose synaptic connections (Coleman & Yao, 2003). The precise role of Aβ in this process remains under investigation.

Neuronal hyperexcitability appears to be an early feature of AD (Targa Dias Anastacio *et al*, 2022). In patients, increased brain activity is observed during cognitive tasks (Celone *et al*, 2006). Preclinical animal and cell models show elevated firing rates and lower activation thresholds (Šišková *et al*, 2014). These alterations, detected across multiple scales from single neurons to entire regions, suggest hyperexcitability may serve as an early biomarker, although further research is needed to confirm this.

Since swallowing difficulties and autonomic nervous system (ANS) dysfunctions are known to occur in AD, it is plausible that the major component of the ANS, the enteric nervous system (ENS), is also involved in the disease’s etiology, like observations in Parkinson’s disease (Cersosimo & Benarroch, 2012).

The ENS shares key features with the central nervous system (CNS), including transcriptomic profiles and neurotransmitter signaling. Amyloid precursor protein, from which Aβ is derived, is naturally expressed by enteric neurons and glial cells (Arai *et al*, 1991; Cabal *et al*, 1995), supporting the potential involvement of the ENS in AD pathophysiology. Disruptions in gut microbiota and the neuroimmune system, along with gastrointestinal (GI) symptoms, have been linked to AD (Niesler *et al*, 2021a; Fung *et al*, 2020). Immunoreactive Aβ plaques have been found in gut tissue of both patients and animal models (Joachim *et al*, 1989; Van Ginneken *et al*, 2011). Dysbiosis, impaired barrier function, and Aβ accumulation in the colon are associated with neuroinflammatory processes via the gut–brain axis and seem to be key contributors to the onset of AD (Liu *et al*, 2023b; Vogt *et al*, 2017). Transgenic mouse models show early GI abnormalities, including microbial imbalance, inflammation, glial activation, and ENS neuron loss (Brandscheid *et al*, 2017; Semar *et al*, 2013a; Puig *et al*, 2015). Given that the gut is the largest immune organ in the body, intestinal neuroinflammation may contribute significantly to AD pathogenesis. However, it remains unclear whether intestinal dysmotility, inflammation, or ENS dysfunction is an early initiating factor. To address this, acute gut models are needed to investigate immediate and short-term effects of Aβ on enteric neurons. This study aims to explore such effects using electrophysiological, functional, and molecular analyses.

## Results

### Survival and apoptosis of ENS cells after short-term Aβ treatment

To determine whether transient short time Aβ treatment (72 hours [h]) affects ENS cell viability or apoptosis, we employed Calcein AM / Annexin V-PE / 7-AAD triple-labeling with flow cytometry to distinguish live (Calcein⁺), early apoptotic (Annexin⁺), and late (7-AAD⁺) apoptotic cells. Calcein AM stains viable cells via intracellular esterification, while 7-AAD enters cells with compromised membranes, staining nuclei red (PerCP). Early apoptosis is marked by phosphatidylserine externalization, detected by Annexin V binding. ENS cells were initially gated in P1 (FSC/SSC, Fig. 1A) to exclude debris. Aβ treatment did not alter cell viability (Fig. 1B) or early/late apoptosis (Fig. 1C, D) across all concentrations tested (10nM to 10µM). To validate this, a CellTiter-Glo® luminescence assay (Promega) was performed measuring ATP as an indicator of metabolic activity (Table EV1). Consistent with calcein data, CellTiter-Glo® showed no reduction in viability after 72h Aβ exposure (Fig. 1E). Interestingly, slight ATP increases at 10nM (+9.12% ±7.25) and 100nM (+15.10% ±6.44) suggesting a potential stimulation. No significant changes occurred at 1–10µM (Δ1µM: +5.06% ±8.10; 2.5µM: +2.67% ±7.89; Δ10µM: +0.46% ±5.96).

**Figure 1:**
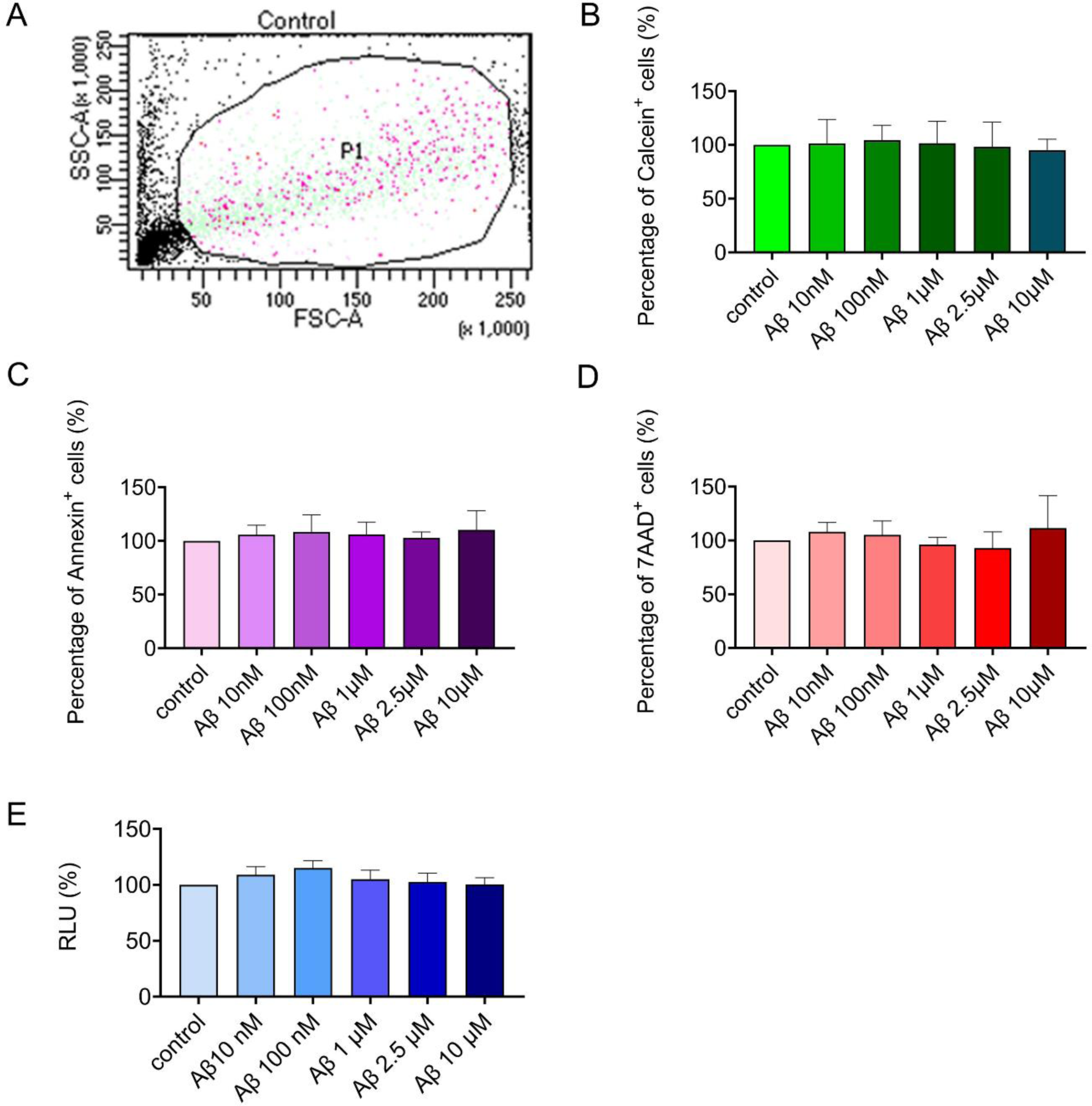
Effect of beta-amyloid (Aβ) on enteric nervous system (ENS) cell survival and apoptosis. To investigate the effects of Aβ on ENS cell viability and apoptosis, primary myenteric plexus cells were isolated from wild-type C57BL/6J mice and treated for 3 days with increasing concentrations of monomeric Aβ (10nM, 100nM, 1µM, 2.5µM, or 10µM, Synpeptide). (A) Flow cytometry (FACS) was used to remove debris and gate viable ENS cells. Viability and apoptosis were assessed in combination via calcein-AM (B), Annexin V (C), and 7-AAD staining (D), all analyzed via flow cytometry. Calcein-positive cells maintained viability, whereas Annexin V and 7-AAD staining revealed no significant increase in early or late apoptotic populations following Aβ treatment. (E) The CellTiter-Glo® luminescence assay further confirmed that metabolic activity was unchanged across the treatment groups. The data represent three independent experiments (n=3 per group) and were normalized to the control. Statistical analysis was performed using one-way ANOVA.

Although short-term Aβ exposure showed no toxic effects on ENS cell survival, subtle, nonlethal responses might still occur. Morphological changes and network alterations, such as neurite outgrowth, often precede cell death. We therefore assessed Aβ’s impact on early stage neuritogenesis.

### Effects of the Aβ peptide on neurite outgrowth

MP was isolated, cultured and challenged with varying Aβ concentrations (10nM– 10µM). Neurite outgrowth was monitored at 4, 24, 48, and 72h using the IncuCyte® Live-Cell Analysis System. Control neurite lengths were normalized to 100% (slope = 0 mm/mm²/h). In general, neurite outgrowth varied depending on Aβ dose and exposure duration (Figs 2 and 3, Tables EV2 and 3; Fig. EV1).

**Figure 2:**
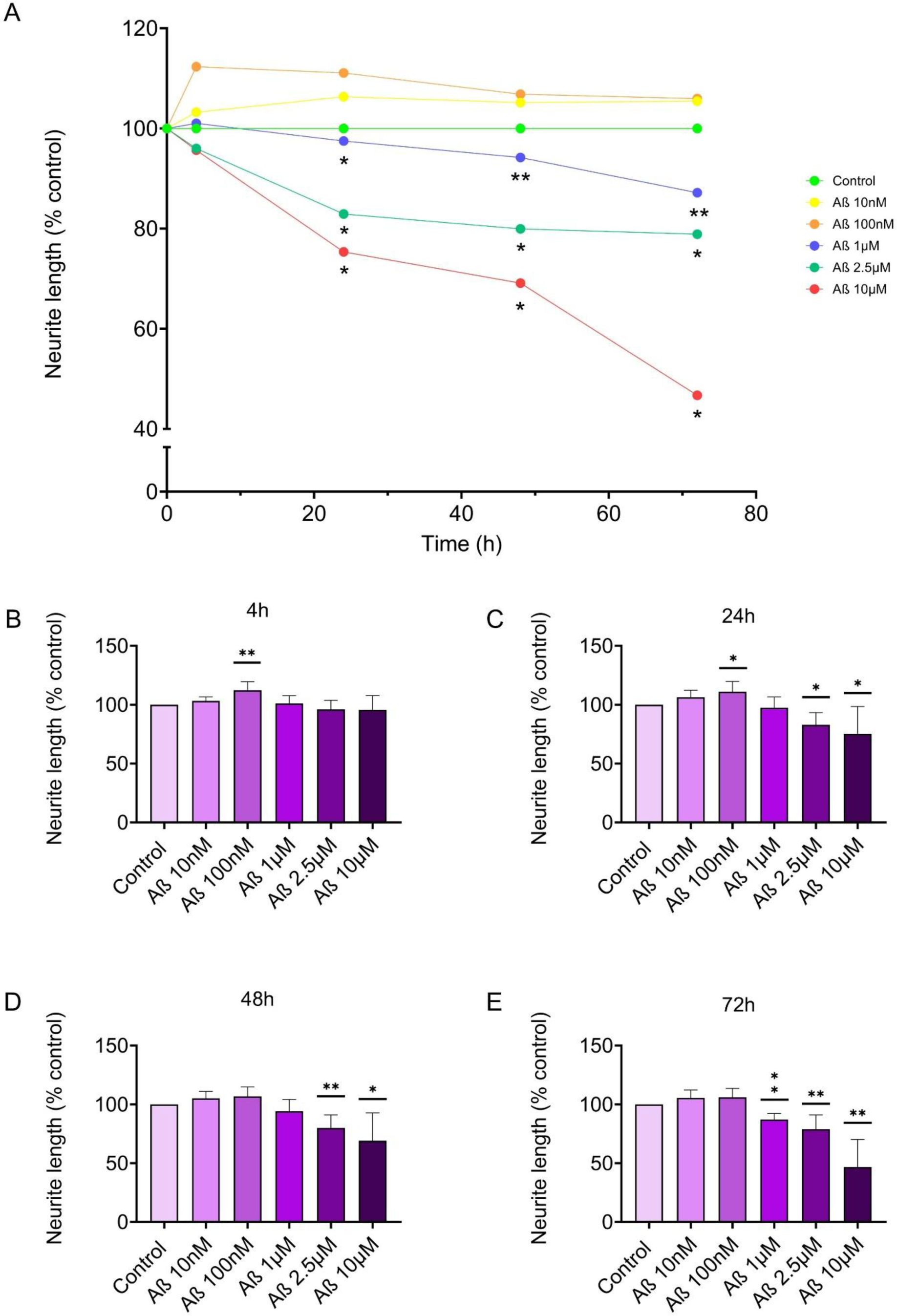
Effect of beta-amyloid (Aβ) on neurite outgrowth in enteric nervous system (ENS) cells. To examine the concentration- and time-dependent effects of Aβ on neurite development, primary ENS cells were treated with increasing concentrations of monomeric Aβ (10nM, 100nM, 1µM, 2.5µM, 10µM), and neurite outgrowth was monitored using the IncuCyte® Live-Cell Analysis System over a 72h period. (A) Course of neurite outgrowth in ENS cells after Aβ exposure for 3 days. Neurite development was progressively impaired in response to increasing Aβ concentrations and longer incubation times. While low nanomolar concentrations had little to no effect or even a transient promoting effect, micromolar concentrations led to a pronounced, time-dependent reduction in neurite length, with the most marked impairment observed at 10µM after 72h. (B) After 4h, co-culturing ENS cells with Aβ had no detrimental impact on neurite lengths, independent of the amyloid concentration used. Exposure of ENS cells to Aβ for 24h (C), 48h (D) and 72h (E) led to a significant reduction in neurite length at concentrations of 1µM, 2.5µM and 10µM. Data are normalized to those of the control and time point 0h. * *p* < 0.01, ** *p* < 0.05 and *** *p* < 0.001 vs. control by one-way ANOVA. Aβ 10nM n=5; Aβ 100nM, 1µM, 2.5µM and 10µM: n=6 per group.

**Figure 3:**
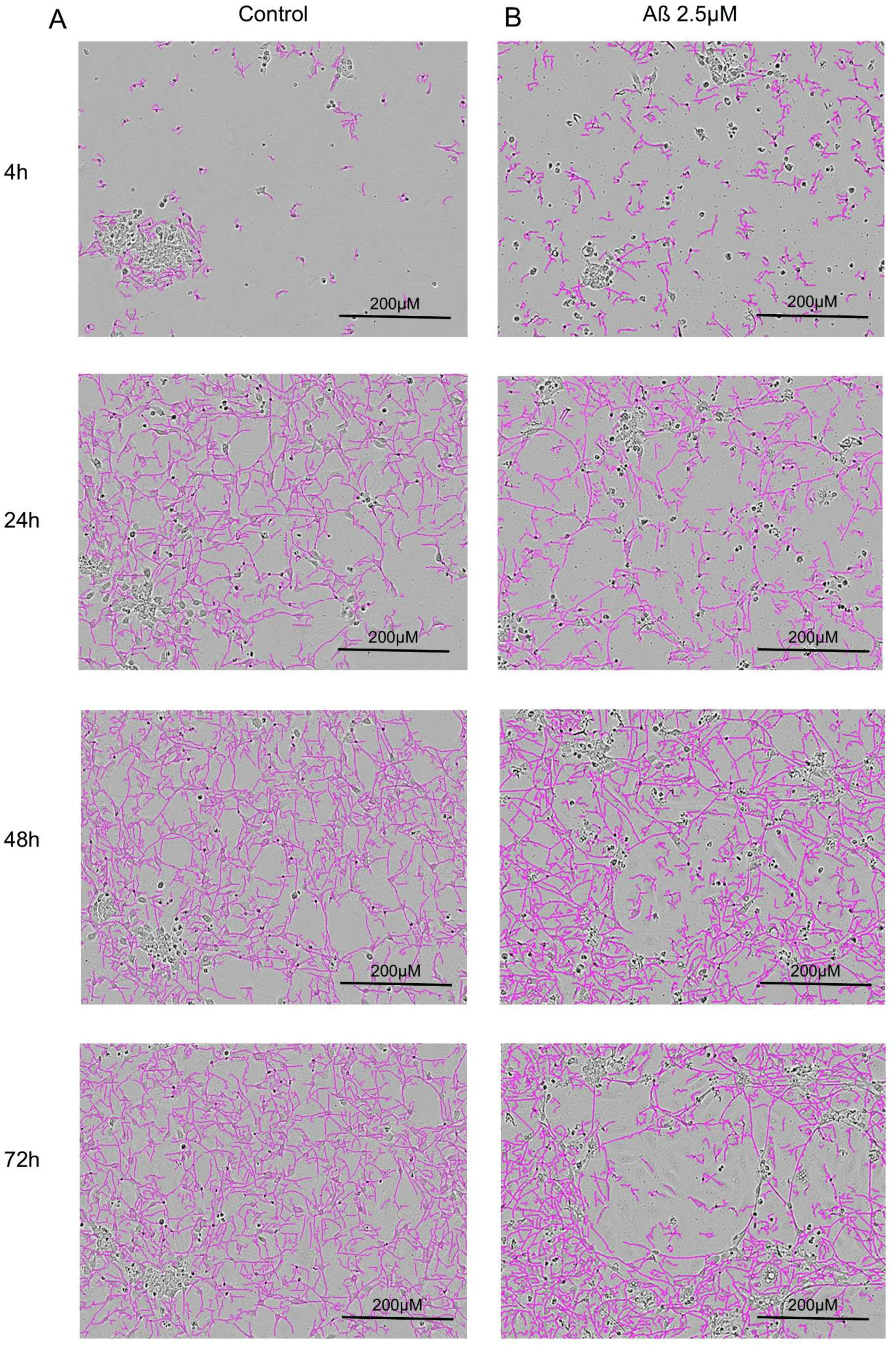
Influence of beta-amyloid (Aβ) treatment for 72 hours (h) on neurite outgrowth in enteric nervous system (ENS) cells. To visualize the Aβ-induced changes in neurite morphology, ENS cells were cultured for 72h with increasing concentrations of monomeric Aβ (10nM, 100nM, 1µM, 2.5µM, 10µM; only representative images of the 2.5µM condition are shown here) and imaged with the IncuCyte® Live Cell Analysis System. (aA) Control cells exhibit extensive neurite networks (purple), as determined by NeuroTrack® software. (B) Treatment with 2.5µM Aβ resulted in a visible reduction in neurite growth and network complexity. The purple overlay shows the neurites identified and quantified by the Incucyte® NeuroTrack® mask, highlighting the differences in neurite length and density between conditions. The images correspond to the endpoint after 72h and are representative of the inhibitory effects quantified in Fig. 2. n=6

For both nanomolar conditions (Figs. 2, 3), we found slight increases for all the time points investigated. Importantly, for 100nM Aβ a clear neurotrophic effect occurred after 4h (+12.33% ± 7.23) and 24h (+11.08% ± 8.74) but then declined mildly after 48h (+6.84% ± 8.01; Tables EV1 and 2). Compared with the control, nanomolar Aβ (10nM and 100nM) still slightly promoted neurite growth after 72h: +5.50% ± 6.78 (10 nM) and +5.97% ± 7.64 (100 nM), as already confirmed by CellTiter-Glo® data (Fig. 1E, Table EV1).

In contrast, micromolar Aβ caused a dose-dependent reduction: at 2.5 µM and 10 µM, neurite length decreased by 17.06% to 53.28% within 24 to 72h (Figs. 2, 3; Tables EV 1 and 2). Furthermore, after 72h of incubation, a significant reduction in neurite length was already observed at an Aβ concentration of 1 µM (-12.83% ± 5.26). Linear regression, including 0h values and calculated growth rates (slopes, mm/mm²/h), is shown in scatter plots with regression lines (Figure EV1). Slopes at 1, 2.5, and 10µM (-0.179, -0.288, -0.697 (mm/mm²/h) were significantly negative (*p* = 0.0005, *p* = 0.0001, *p* = 0.0001), unlike those at 10nM and 100nM (0.057, 0.0003; *p* = 0.181, *p* = 0.995).

These results suggest that low (nanomolar) Aβ concentrations slightly promote neurite growth, while relatively higher (micromolar) levels reduce neurite length in a dose- and time-dependent manner, highlighting the dual effect of Aβ on neurons. Such a biphasic response may indicate a contribution to neuronal hyperexcitability, with low levels transiently enhancing activity, while higher concentrations disrupt homeostasis and compromise neurite integrity. Notably, neurite length did not decrease further even after 120 and 140h of Aβ exposure (data not shown).

To confirm these findings, we conducted immunocytochemical class III β-tubulin 1 (Tuj1) stainings after 72h to visualize neurite structure. Consistent with the IncuCyte data, Tuj1 staining showed a significant reduction in total neurite outgrowth and signal intensity from 1µM Aβ upward (1µM: –27.50%±5.31; 2.5µM: –42.99% ±13.91; 10µM: –51.38% ±13.92; Fig. 4D-F; Fig. 5A; Table EV4) compared with the control (Fig. 4A), while nanomolar doses showed no impairment (10nM: –5.91% ±2.65; 100nM: – 3.16% ±5.56; Fig. 4A-C; Fig. 5A; Table EV4) compared with the control (Fig. 4A). Of note, total cell number was not affected, shown by 4′,6-diamidino-2-phenylindole (DAPI) staining. These results reinforce a concentration-dependent vulnerability, where only high Aβ levels compromise structural integrity.

**Figure 4:**
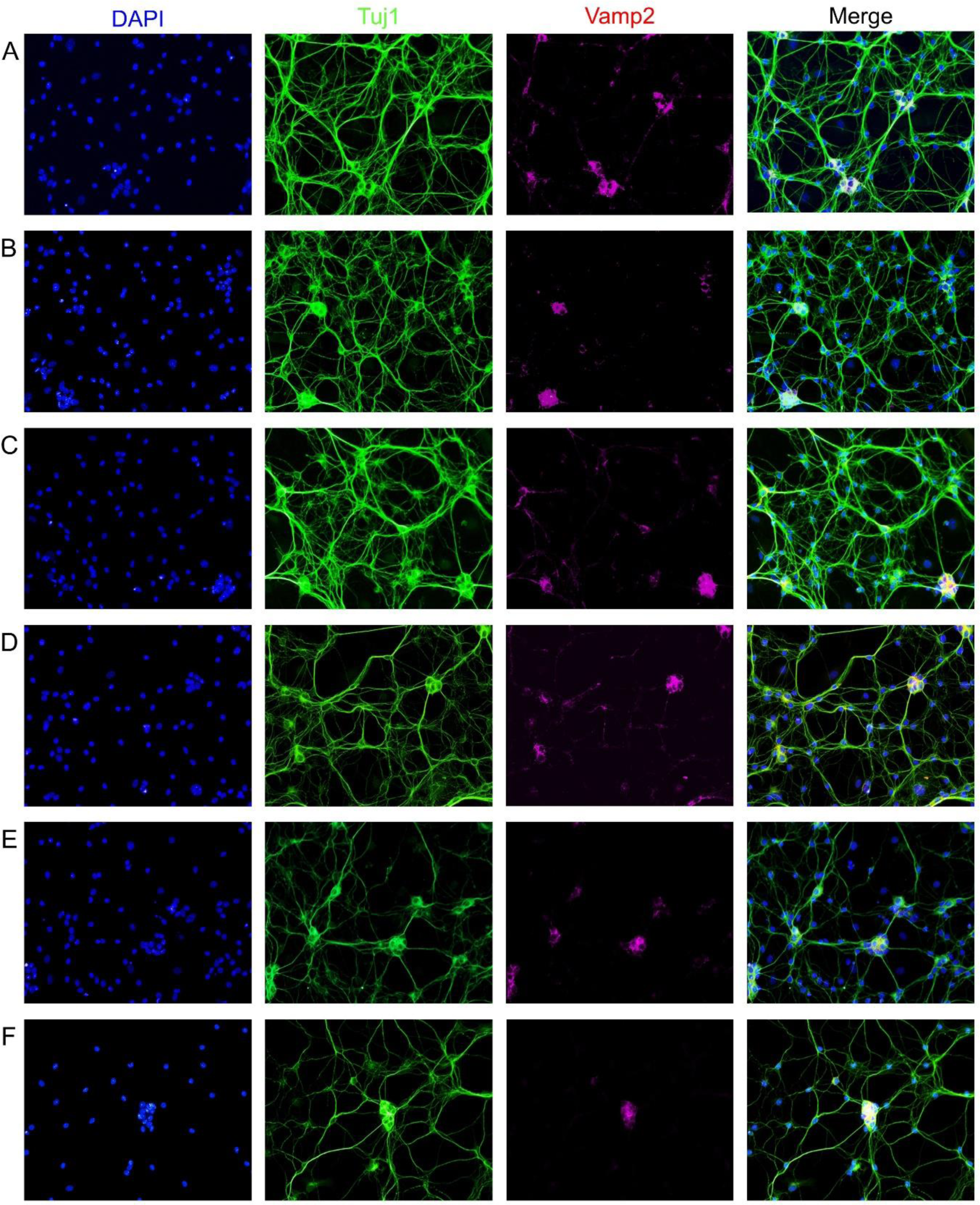
Immunocytochemical staining of enteric nervous system (ENS) cells incubated for 72 hours (h) together with beta-amyloid (Aβ). ENS cells were isolated and cultured together with Aβ for 72h. (A) Control, (B) Aβ 10nM, (C) Aβ 100nM, (D) 1µM, (E) 2.5µM, (F) Aβ 10µM. Nuclei are visualized by 4′,6-diamidino-2-phenylindole (DAPI, blue). The neuronal structures were visualized by immunostaining for neuron-specific class III β-tubulin (Tuj1, green), which revealed the entire neurite network and structural integrity. Synapse density is shown by vesicle-associated membrane protein 2 (Vamp2) staining in magenta. The merged images represent the overlay of all channels, highlighting the spatial distribution and potential colocalization of the proteins. n=3 per group

**Fig. 5.**
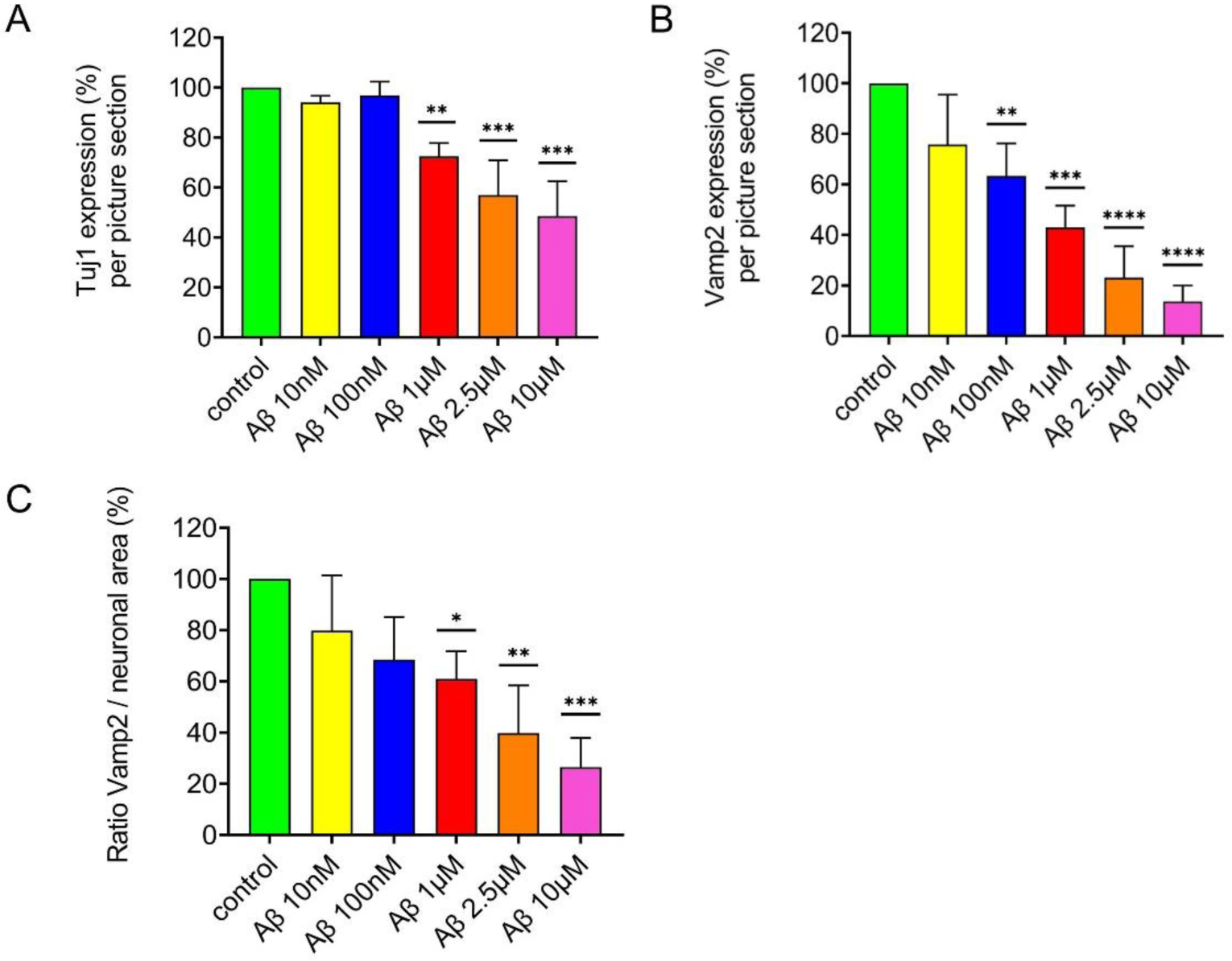
Quantification of the immunocytochemical staining of enteric nervous system (ENS) cells incubated for 72 hours (h) together with beta-amyloid (Aβ). (A) Exposure to micromolar concentrations of Aβ resulted in a significant reduction in neuron-specific class III β-tubulin (Tuj1) expressions and density, whereas nanomolar concentrations did not significantly affect neurite morphology). (B) The synaptic sites were labeled with vesicle-associated membrane protein 2 (Vamp2, red), whereas synaptic density was already reduced at nanomolar Aβ concentrations. This synaptic loss became increasingly pronounced with increasing Aβ concentrations). (C) The ratio of Vamp2 to Tuj1 signals was used to distinguish synaptic loss from general degeneration of neurites. A significant decrease in this ratio at micromolar Aβ concentrations suggests that synapses are disproportionately affected compared with the overall neuronal structure. The data are normalized to the control. * *p* < 0.01, * *p* < 0.05 and ** *p* < 0.001 vs. control by one-way ANOVA. n=3 per group

### Aβ-induced synapse loss and damage in the ENS

We next examined whether neurite degeneration corresponds with synaptic changes, using Vamp2 staining to quantify presynaptic density after short-term exposure(Yan *et al*, 2022). Vamp2 staining enabled simultaneous analysis of neuronal areas and synapse number. Notably, compared with the control (Fig. 4A), even nanomolar Aβ reduced synaptic density: 10nM induced a mild (–24.16% ±19.78; Fig. 4B; Fig. 5B; H; Table EV5) and 100nM a significant (–36.60% ±12.92; Fig. 4C, Fig. 5B) reduction.

To distinguish synapse loss from neurite atrophy, we calculated the Vamp2/Tuj1 ratio, which significantly declined only at ≥1µM Aβ (Fig. 5C, Table EV6), confirming specific synaptic vulnerability. Thus, Aβ reduces synaptic density in a dose-dependent manner, with effects emerging already at nanomolar levels. These changes reflect direct synaptic sensitivity rather than secondary loss due to neurite shrinkage, highlighting Aβ’s functional impact on enteric connectivity.

### Aβ induces Hyperexcitation in Enteric Neurons

To test whether Aβ affects ENS network activity, we performed electrophysiological recordings using microelectrode arrays (MEAs). Within 3–5 days *in vitro*, cultured MP developed spontaneous activity that could be recorded via single-chip MEAs (60MEA200/30iR-Ti, MCS Reutlingen, Fig. 6A, B). Across 17 independent WT-derived cultures, activity was validated pharmacologically, by reduction of neuronal firing using the sodium channel blocker tetrodotoxin and conversely increasing the number of spikes through the addition of the neurotransmitter acetylcholine (100µM). Electrode activity remained stable over time. Following a 10-minute baseline, Aβ (10nM–1µM) was acutely applied, and spike rates recorded. While 10nM did not increase the baseline activity (109.42 ±33.69 spikes/min) compared with the control (115.67±29.40 spikes/min), 100nM increased firing to 203.48 ± 45.78 spikes/min (+76%), and 1µM reached 193.42±45.27 spikes/min (+67%) (Fig. 6C), confirming rapid Aβ-induced hyperexcitation. Higher concentrations were omitted as effects plateaued. Representative raster plots (10 sec) illustrate the acute increase in spike activity in a culture exposed to 100nM Aβ, with a clear increase in firing immediately after administration (Fig. 6D-E). Thus, Aβ rapidly enhances firing in ENS neurons, reflecting functional excitability changes beyond structural damage.

**Figure 6:**
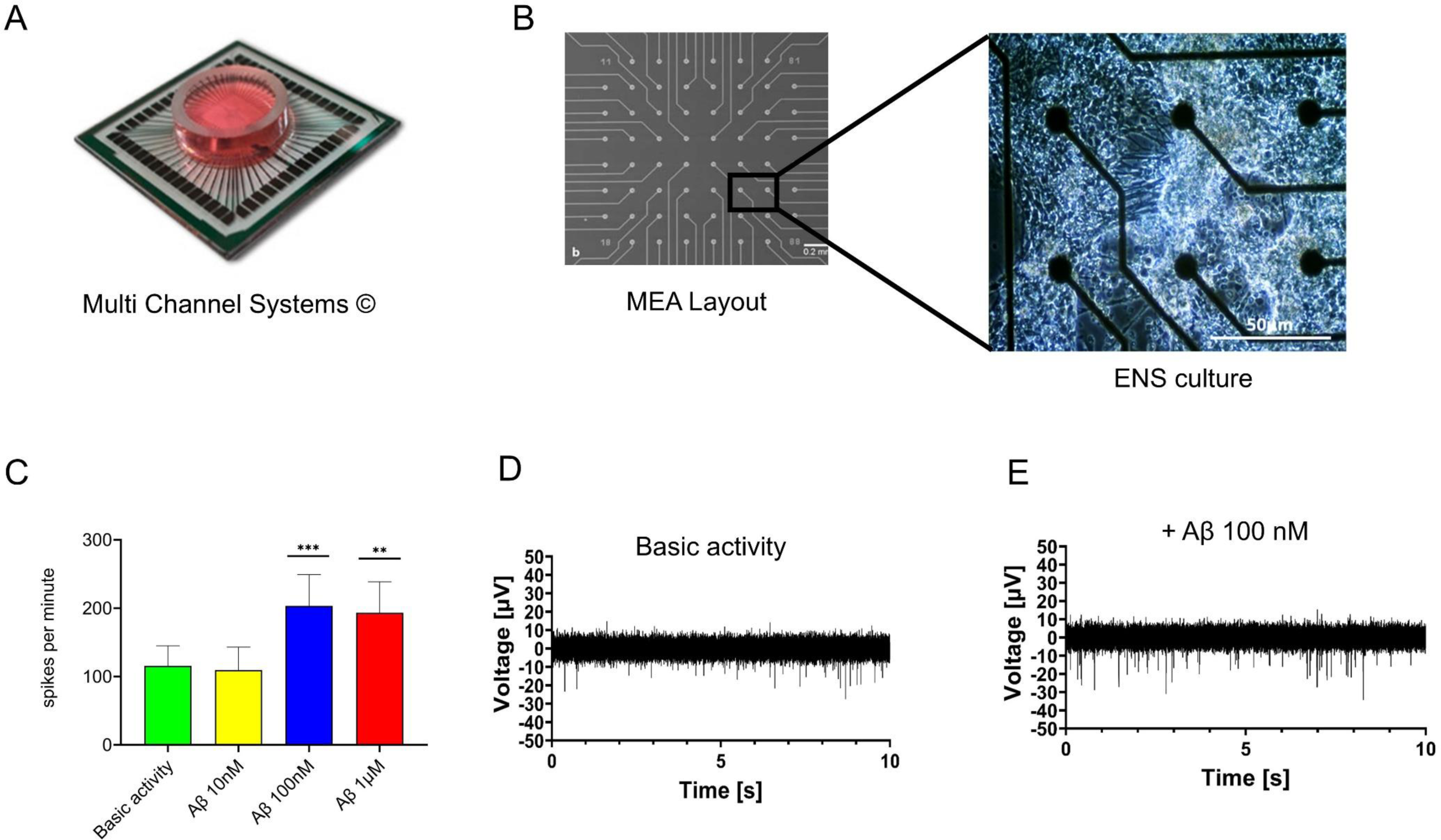
Microelectrode array (MEA) recordings of myenteric plexus cells. (A) Illustration of the planar MEA chip (60MEA200/30iR-Ti, Multi-Channel Systems) with 60-electrode architecture (B) used for extracellular recordings from cultured enteric nervous system (ENS) cells grown on the MEA chip showing a network of enteric neurons and glial cells suitable for electrophysiological analysis. (C) Spontaneous neuronal activity was recorded for 10min prior to acute administration of various Aβ concentrations (10nM, 100nM, 1µM). Subsequent 10min-recordings revealed a clear increase in spike frequency. While 10nM Aβ had no significant effect on the neuronal firing rate, both 100nM and 1µM induced a marked and statistically significant increase in spike activity, demonstrating acute hyperexcitability of ENS neurons in response to Aβ (D,E). Representative raster plots and spike rate histograms from ENS cultures treated with 100nM Aβ illustrate the rapid and robust increase in the neuronal firing rate immediately following Aβ exposure. This acute excitatory effect suggests that Aβ not only impairs neuronal structure but also directly modulates the functional activity of the neuronal network. ** *p* ≤ 0.01, *** *p* ≤ 0.001 using a one-way ANOVA. The sample sizes ranged from 3 to 6 independent cultures per condition.

### Aβ increases calcium influx in ENS cells

To further investigate the underlying mechanisms of this response, we assessed intracellular calcium using fluo-4-based imaging. Action potentials and synaptic activity are closely linked to Ca²⁺ influx via voltage-gated channels and N-methyl-D-aspartate (NMDA) receptors. Neurons were identified by morphology and ganglionic localization (Fig. 7A). Baseline recordings were followed by acute Aβ exposure (10nM–1µM) and Ca²⁺ tracking over 10min. Cell viability was verified via high-potassium depolarization post-recording (not shown). Fluorescence intensity changes (ΔF/F) were visualized via a false-color scale, with red indicating increased intracellular calcium (neuronal activation) and blue reflecting decreased activity (inhibition) (Fig. 7A). Under baseline conditions, ENS neurons exhibited low but detectable spontaneous Ca²⁺ transients (Fig. 7B; Videos EV1 and 3). Aβ triggered immediate, significant increases in intracellular Ca²⁺ at all concentrations. The response plateaued beyond 10nM. Spatially, nanomolar Aβ mainly increased Ca²⁺ influx in neurites (presynaptic sites), while micromolar doses affected somata (Videos EV2 and 4), possibly reflecting excitotoxic stress. These findings confirm rapid Aβ-induced excitability changes at the single-cell level, supporting MEA data and suggesting early functional disruption of enteric neurons.

**Figure 7:**
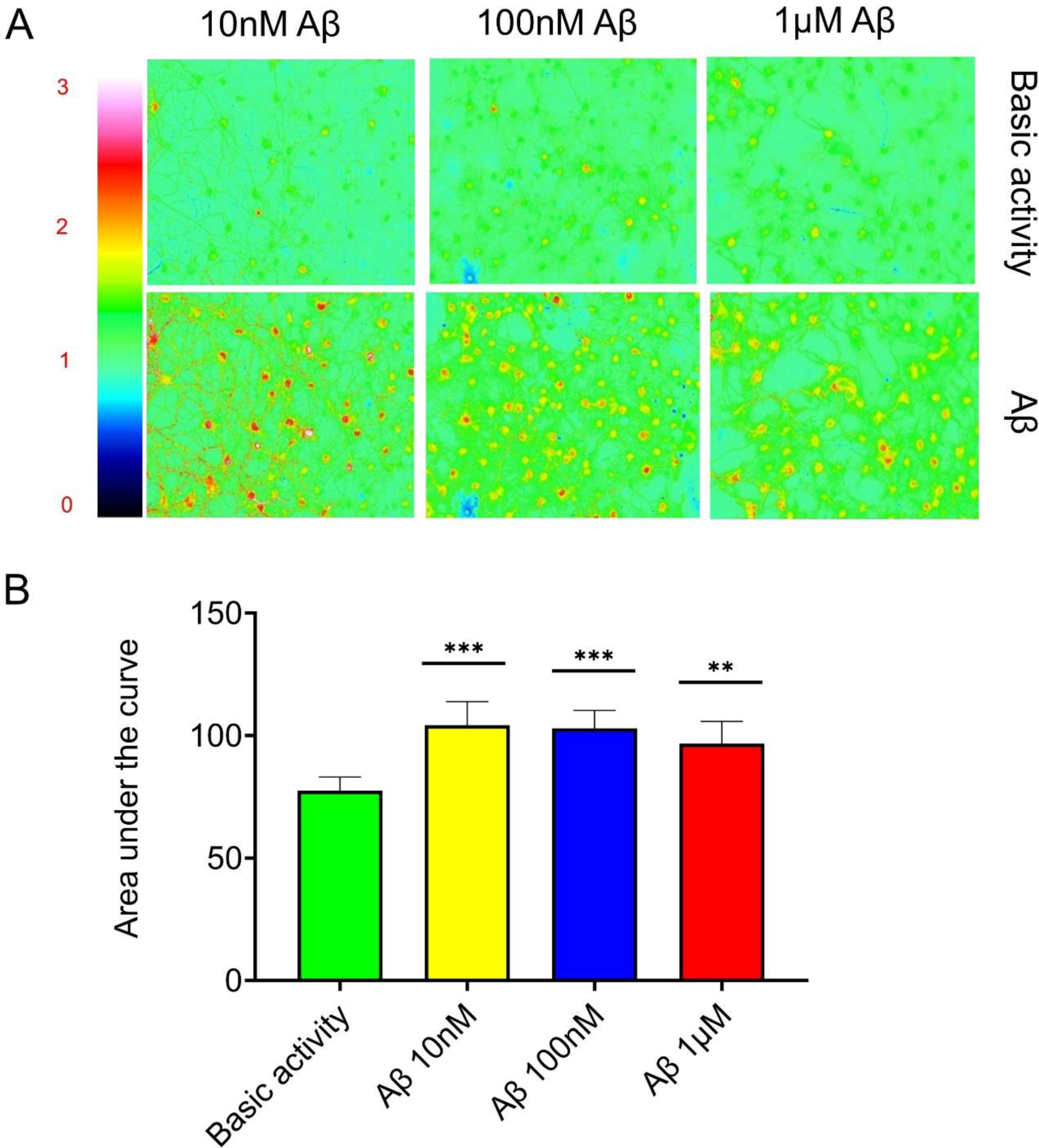
Calcium imaging analysis of Aβ-induced neuronal activation in enteric nervous system (ENS) cells. (A) False-color-coded fluorescence images showing the intracellular Ca²⁺ levels in cultured ENS cells loaded with the Ca²⁺-sensitive dye Fluo-4. Yellow to red tones indicate increased intracellular calcium concentrations (neuronal activation), whereas blue tones reflect decreased activity or inhibition. The neurons were identified on the basis of their characteristic morphology, their localization within the ganglia and their dye loading. Spontaneous Ca²⁺ transients are visible prior to treatment. Acute administration of Aβ at concentrations of 10nM, 100nM, and 1µM triggered a rapid and sustained increase in intracellular Ca²⁺ levels, with spatial differences in activation patterns observed: nanomolar Aβ preferentially increased Ca²⁺ influx in neurites, whereas micromolar Aβ predominantly increased somatic calcium. (B) Ca²⁺ dynamics were quantified by calculating the area under the curve (AUC) of fluorescence intensity changes (ΔF/F) over time. Compared with baseline, Aβ treatment resulted in a statistically significant increase in intracellular Ca²⁺, with a plateau effect observed between 100nM and 1µM, suggesting maximal neuronal activation at lower nanomolar doses. * *p* ≤ 0.05, ** *p* ≤ 0.01, *** *p* ≤ 0.001 via one-way ANOVA. Aβ 10nM and 100nM: n=5; Aβ 1µM: n=6 per group

### Combined luminal and mesenteric ex vivo perfusion experiments

To assess physiological outcomes of ENS network changes, we analyzed peristalsis in *ex vivo*-mesenteric-perfused jejunal segments from WT mice(Schreiber *et al*, 2014; Gries *et al*, 2021). Movements were recorded in real time following luminal Tyrode’s buffer (baseline) and arterial perfusion with 100nM Aβ (Fig. 8A-C). Video analysis (MotMap, LabView) quantified mean interval (reciprocal of contraction frequency), duration of dilation and contraction phases, and amplitude of contractions (Δ-amplitude), which represents the strength and extent of smooth muscle contractions. Aβ had no effect on contraction frequency or duration (Fig. 8D-F; Videos EV 5 and 6), but significantly reduced contraction amplitude (Fig. 8G-I), indicating impaired neuromuscular coupling or increased excitatory neurotransmission under acute Aβ influence by impairing the contractile force.

**Figure 8:**
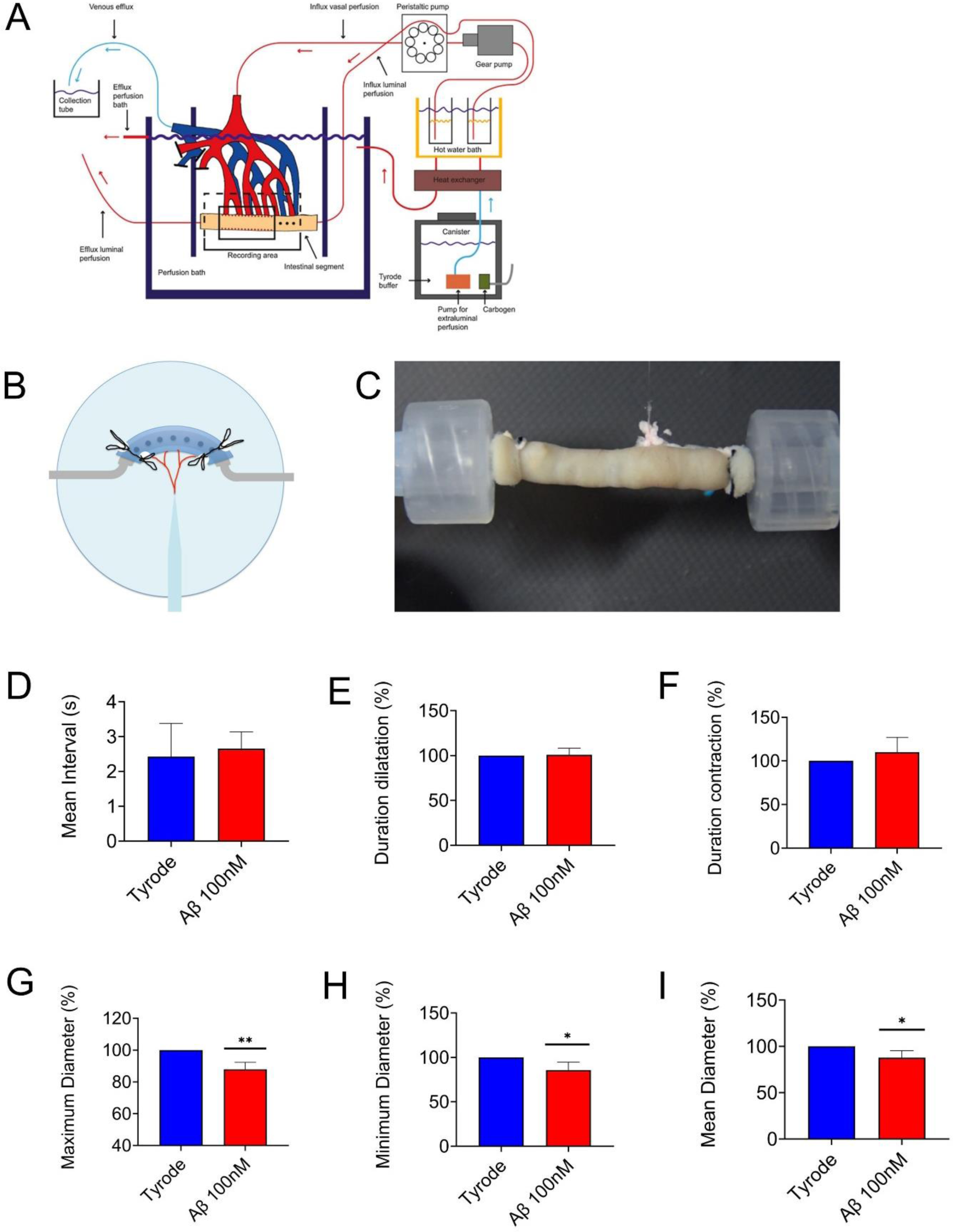
Analysis of intestinal motility during combined ex vivo luminal mesenteric perfusion experiments on wild-type C57BL/6J mice. (A) Experimental setup illustrating the combined mesenteric artery and lumen perfusion of a 2.5 cm long jejunal segment (Schreiber *et al*, 2014) mounted in an organ bath filled with Tyrode’s buffer (B and C). Beta-amyloid (Aβ, 100nM) was administered via mesenteric artery cannulation using a syringe pump to simulate systemic exposure. Intestinal movements were continuously recorded on video and quantitatively analyzed via MotMap and LabView software to evaluate motility parameters. (D) The mean interval between contractions, which represents the reciprocal of the contraction frequency, was measured before and during Aβ perfusion. No significant changes were observed, i.e., the timing of peristaltic contractions remained stable under acute Aβ exposure. Similarly, the duration of the dilation phase (relaxation, E) and the contraction phase (F) of the intestinal segments did not significantly change after Aβ treatment. (G-I) In contrast, the contraction amplitude, a measure of the strength and extent of smooth muscle contractions, was significantly reduced during Aβ perfusion. This decrease was evident in both the mean contraction amplitude and the minimum and maximum diameters of the jejunal segments, indicating impaired contractile force. * *p* < 0,05; ** *p* < 0,01 and *** *p* < 0,001 vs. Tyrode’s test by Student’s t test. n=5 animals

Within the ENS, rapid excitatory neurotransmission between interneurons and motor neurons is known to be mediated by nicotinic acetylcholine receptors (nAChRs), facilitating the coordinated smooth muscle contractions necessary for peristalsis and other motility patterns(Rueda Ruzafa *et al*, 2021). While nAChR antagonists can suppress ENS-mediated motility responses, nicotine and other nAChR agonists can stimulate intestinal motility via this excitatory pathway(Gershon, 1999). To explore their involvement in the context of acute AD, and to determine whether Aβ also acts as an agonist of nAChRs, rat intestines mesenterically with Aβ, the agonist nicotine, and methyllycaconitine (MLA), a specific inhibitor of the α7 subunit of the nAChR, along with Aβ. Notably, arterial administration of nanomolar Aβ-containing conditioned medium to Aβ_42_-transfected SH-SY5Y cells (1.4ng/ml, not shown) was used in this sub experiment to simulate a more physiological environment with biologically produced Aβ_1-42_. Administration of the agonist nicotine led to a significant, dose-dependent disruption of contractility (Fig. EV2A). Similar to nicotine alone, the administration of Aβ significantly impaired motility and led to a marked increase in contractility comparable to that observed with nicotine (Fig. EV2B). To demonstrate that the Aβ-induced motility change is nAChR-mediated, we perfused the intestine not only with Aβ but also in combination with the inhibitor of the α7 subunit MLA. MLA administration significantly reversed the Aβ effect (Fig. EV2C).

Although short-term Aβ exposure does not induce neuronal death, it triggers structural and functional alterations in ENS neurons at nanomolar concentrations. These include reduced neurite and synapse integrity, increased excitability, elevated Ca²⁺ influx, and impaired intestinal contractility. Together, these findings suggest that the ENS is an early and sensitive target of Aβ-related pathology.

## Discussion

AD is a debilitating neurodegenerative disease characterized by Aβ deposition and tau hyperphosphorylation(Wang *et al*, 2017). Research has primarily focused on CNS lesions, particularly in the hippocampus and frontal cortex, that lead to neuronal death. However, increasing evidence suggests that early peripheral abnormalities, including GI symptoms like inflammation, swallowing dysfunction and constipation, as well as fecal incontinence, also occur in AD patients(Wang *et al*, 2017; Niesler *et al*, 2021b), whereby intestinal activity is regulated by the ENS(Rao & Gershon, 2016). Nevertheless, research focusing upon pathological features in the GI tract or the ENS of AD patients is only slowly increasing and still very limited.

Aβ deposition has been identified not only in the CNS but also in the peripheral nervous system, suggesting local Aβ production(Joachim *et al*, 1989). Given that AD pathology develops decades before clinical symptoms emerge, early events likely occur during the preclinical phase. Studies in aged mice show Aβ accumulation in the ENS, leading to inflammation, motility disturbances, and neuronal loss(Puig *et al*, 2015). One longitudinal study also identified severe, early-stage ENS impairment preceding CNS pathology. These changes appear to follow a temporal pattern, with early increases in inflammatory and neural stem cell markers, potentially indicating an initial phase of functional decline(Semar *et al*, 2013b).

We believe that these early and severe changes reflect the acute phase of the disease, during which pathological processes are particularly pronounced. Our study focused on acute and short-term effects of monomeric Aβ on the ENS and GI motility. We hypothesized that Aβ triggers neuronal overexcitement, which ultimately leads to exhaustion, a kind of “burnout,” and thus to dysfunction or cell loss over time. First, we assessed viability after Aβ exposure; second, we analyzed neuritic and synaptic integrity; and third, we evaluated electrophysiological and functional responses. While short-term monomeric Aβ exposure did not induce apoptosis or reduce viability, high micromolar concentrations reduced neurite outgrowth and neuronal area, indicating early structural changes without cell death. This aligns with Neniskyte et al.(Neniskyte *et al*, 2011), who found that monomeric Aβ_1-42_ causes neuronal loss at sub-micromolar levels without direct toxicity. These authors proposed two mechanisms of monomeric Aβ toxicity: (1) direct neurotoxicity at high concentrations, and (2) indirect microglia-dependent toxicity at lower levels, requiring longer exposure.

Our results are consistent with studies on hippocampal cultures(Cappai & Barnham, 2008), as given the physiological Aβ concentrations in the brain (soluble: 2–65 nM; insoluble: 200–4500nM (Kuo *et al*, 1996; Hellström-Lindahl *et al*, 2009), the pathophysiological relevance of direct Aβ toxicity *in vivo* is questionable, especially since >100 µM may be required(Liao *et al*, 2007). Thus, nanomolar-range indirect effects appear more relevant(Brown & Neher, 2010).

Importantly, the aggregation state of Aβ critically affects toxicity(Cizas *et al*, 2010). Monomers transition into oligomers via α-helix to β-sheet conversion, then form protofibrils and insoluble fibrils, which is the basis of plaques(Imbimbo *et al*, 2023; Penke *et al*, 2020b). Aggregation begins above a monomer threshold of ∼90nM(Novo *et al*, 2018). Thus, it is not surprising that, in our study, nanomolar Aβ had no effect on cell death or neurite growth, but synaptic structure and function were impaired at these levels, highlighted by MEA recordings, calcium imaging, and motility assays. This indicates that Aβ affects neuronal function at low concentrations but causes structural damage only at higher levels.

The mechanism behind Aβ-induced enteric neurite inhibition remains unclear. In the CNS, Aβ disrupts cytoskeletal dynamics and reduces neurotrophins like brain-derived neurotrophic factor (BDNF) and nerve growth factor (NGF)(Matrone *et al*, 2008). Furthermore, Aβ reduces the expression of neurotrophins such as BDNF and NGF, which are essential for neurite outgrowth(Matrone *et al*, 2008). BDNF, also expressed in the ENS(Boesmans *et al*, 2008), supports neurite growth and plasticity. BDNF deficiency has been linked to GI dysfunctions such as Hirschsprung’s disease(Chalazonitis, 2004). In animal models, BDNF blockade has been shown to lead to shortened neurite length and reduced neuronal connectivity in the ENS(Liu, 2018). Likewise, NGF deficiency impairs enteric neurite growth and causes motility issues(Wong *et al*, 2019). Thus, impaired neurotrophin signaling or macrophage-mediated toxicity may contribute to reduced neurite growth after Aβ exposure. Here, further research is needed.

In addition to neuronal loss, synaptic degeneration is a hallmark of AD. Aβ oligomer levels in the brain correlate with cognitive impairment and loss of synaptic markers like Vamp2( Pham et al, 2010). Consistently, we found that Vamp2 expression in the ENS decreases with Aβ exposure, even at nanomolar levels. Notably, this reduction was independent of total neuronal area, as shown by a decreased Vamp2/Tuj1 ratio at ≥1µM. Since Vamp2 is vital for vesicle fusion and synaptic transmission, its downregulation may impair communication and explain synaptic dysfunction in AD. Aβ can disrupt Vamp2–SNAP-25 interactions, impairing SNARE complex formation and neurotransmitter release(Sharda *et al*, 2020). Our data suggest Vamp2 downregulation may harm enteric neuronal communication, contributing to GI symptoms.

Moreover, it was recently shown that the administration of Aβ to hippocampal neurons induces Aβ internalization(Russell *et al*, 2012). While electrophysiological Aβ effects in the CNS have been studied intensely(Schulte *et al*, 2021, 2024), little is known about these effects in the ENS. We observed that acute Aβ exposure significantly increased spontaneous spike activity in enteric neurons, suggesting that Aβ can rapidly modulate the excitability of enteric networks. These findings are consistent with CNS reports of increased firing and disrupted network synchronization upon Aβ exposure(Kellner *et al*, 2014) and that neuronal cultures respond to aggregated forms of Aβ with increased spike frequency and impaired long-term potentiation(Schulte *et al*, 2021). Whether hyperactivity or silencing is the initial trigger remains unclear (Busche *et al*, 2008). Nonetheless, our study is the first to confirm direct Aβ-induced hyperexcitability in the ENS.

Calcium imaging supported these findings, showing rapid intracellular Ca²⁺ increases upon Aβ application, in line with studies in WT (Busche *et al*, 2012). Aβ likely induces excitotoxicity by disrupting glutamate balance and overstimulating NMDA receptors, leading to oxidative stress and neuronal dysfunction(Talantova *et al*, 2013). Aβ also enhances long-term depression (LTD), reducing glutamate reuptake and further activating NMDA receptors (Li *et al*, 2009).

Another key finding was the Aβ-induced alteration in intestinal motility, potentially mediated via nAChRs. These receptors regulate GI motility and are present on enteric neurons. CNS studies have shown Aβ binds α7-nAChRs with high affinity(Wang *et al*, 2000), enhancing excitability and upregulating receptor expression(Liu *et al*, 2013). Blocking these receptors prevents Aβ-induced hyperactivity, indicating α7-nAChRs as mediators of Aβ action. The observed increase in contractility following perfusion with Aβ_1-42_-conditioned medium indicates that Aβ might act as an agonist of nAChRs at nanomolar concentrations, suggesting direct activation of excitatory cholinergic networks in the ENS. Crucially, simultaneous perfusion of Aβ with MLA, a selective antagonist of the α7 nAChR subunit, significantly reversed the Aβ-induced changes in motility. These findings support the hypothesis that the effect of Aβ is mediated primarily via the α7 subunit of nAChRs. These results imply that Aβ not only plays a role in the pathophysiology of AD but also may be involved in the modulation of peripheral cholinergic systems. Previous CNS studies have also shown that Aβ can bind with high affinity to nAChRs, particularly the α7 nAChR subtype, leading to increased neuronal excitability in cultures of murine hippocampal neurons and posttranslational upregulation of the α7 subunit of nAChRs. Blockade of nAChRs *in vitro* prevents both increased neuronal excitability and changes in receptor function, indicating a central role of α7-nAChRs in mediating Aβ-induced neuronal hyperexcitation (Liu *et al*, 2013).

As already illustrated by the MEA and Ca imaging experiments, we also demonstrate in our motility assays that the ENS is affected by Aβ by significant changes at nanomolar, i.e., physiological, concentrations. Other studies also show that even low Aβ levels activate nAChRs(Liu *et al*, 2001; Dineley *et al*, 2002). We propose that Aβ binds to and modulates nAChRs in the ENS, impairing motility. This aligns with transgenic mouse models showing early gut Aβ deposition, impaired gastric emptying, and reduced cholinergic/nitrergic neurons (Liu *et al*, 2023a; Semar *et al*, 2013a). Our Tuj1 data support this neuronal loss(Sun *et al*, 2020) after Aβ exposure. Moreover, direct Aβ injection into the stomach wall increases jejunal contraction rates(Sun *et al*, 2020).

Our data provide evidence that Aβ-induced hyperexcitation might be an early stage of AD, which leads, after longer periods to functional and morphological alterations in the enteric neurons and the gut. Even monomeric Aβ disrupts neurite and synaptic structure, increases excitability, and impairs motility, highlighting early gut involvement in AD. These effects are likely mediated via NMDA and nAChRs.

## Methods

### Animals

Newborn C57BL/6J mice (P1‒P4) were used for *in vitro* experiments, while adult wild-type (WT) C57BL/6J mice aged 8 ‒10 weeks and adult Wistar rats of either gender were used for *ex vivo* studies. The animals were housed under specific pathogen-free (SPF) conditions at 22±3°C, with 50–70% relative humidity and a 12h light/dark cycle, in compliance with German regulations. All animals were euthanized in accordance with local ethics committee guidelines and Rhineland-Palatinate animal protection laws. Newborn mice were decapitated, adult mice were euthanized by an overdose of isoflurane, and rats were euthanized by CO₂ asphyxiation.

### Primary neuronal cell culture

The entire intestine was removed and placed in cold MEM-4-(2-hydroxyethyl)-1-piperazineethanesulfonic acid (HEPES) buffer (Thermo Fisher Scientific). Under stereomicroscope control, the intestines were opened along the mesenteric line and rinsed with MEM-HEPES (Thermo Fisher Scientific), and the muscle layer was carefully separated from the submucosal/mucosal layers via watchmaker forceps. The muscle layers were stored in ice-cold MEM-HEPES (Thermo Fisher Scientific) for further processing. To isolate the myenteric plexus (MP), the obtained muscle strips were digested with 0.375mg/ml Liberase (Sigma-Aldrich) and 0.2mg/ml DNase (Sigma-Aldrich) in Hanks balanced salt solution for 2.5h at 37°C as previously described(Grundmann *et al*, 2015). The myenteric networks were then detached from smooth muscle cells via gentle mechanical vortexing/pipetting, collected, and used for further experiments. For the cell culture experiments, the neuronal networks were dissociated either mechanically using pipettes (ganglia) or enzymatically via incubation with Accutase (PanBiotech) for 2min at room temperature. The cells were then cultured either as (PCs) or differentiation cultures (DCs). For PC, the cells were suspended in proliferation medium (1% Bovine serum albumin [BSA], 2% B27 [LifeTechnologies], 0.1% β-mercaptoethanol [Thermo Fisher Scientific], 1% penicillin/streptomycin [VWR], and 10ng/ml mouse epidermal growth factor [Immunotools] and 20ng/ml fibroblast growth factor [Immunotools] in DMEM-F12 [Thermo Fisher Scientific]) and cultured for 1-2 days. For DC, the cells were plated in differentiation medium (1% BSA [Sigma Aldrich], 2% B27 with retinoic acid [Life Technologies], 0.1% β-mercaptoethanol [Thermo Fisher Scientific], 1% penicillin/streptomycin [VWR] and 10ng/ml human glial cell line-derived neurotrophic factor [Immunotools] in DMEM-F12 [Thermo Fisher Scientific]) for up to 3-5 days. For all cell culture experiments with Aβ, the medium was replaced with phenol red-free medium containing B27 without antioxidants (LifeTechnologies) and without β-mercaptoethanol (Thermo Fisher Scientifc) before the addition of Aβ to avoid uncontrolled aggregation of the peptide.

### Synthetic Aβ_1-42_ preparation

Monomeric Aβ_1-42_ peptides (Synpeptide), hereinafter referred to as Aβ, were dissolved at 30μM in C1 assay buffer (130mM NaCl, 10mM HEPES, 5mM KCl, 2mM CaCl_2_, [pH 7.2, all Thermo Fisher Scientfic]). The peptides were fully solubilized via 10min of sonication at room temperature. Any precipitates were removed by centrifugation, and the protein concentrations were reverified. The dissolved peptides were then aliquoted for single use, stored immediately at -80°C and used within 4 weeks, as longer storage would compromise the results. All the experiments independently confirmed that C1 buffer alone had no effect on the results (data not shown).

### Plasmids

The spA4ct-DA variants of the C99 coding sequence were inserted into the pCEP4 expression vector (Invitrogen), resulting in the following constructs: pCEP4-spA4ct-DA-WT, pCEP4-spA4ct-DA-I45F, and pCEP4-spA4ct-DA-V50F. These plasmids have been characterized in earlier publications (Grimm *et al*, 2003).

### Cell line culture and transfections

SH-SY5Y human neuroblastoma cells (Uhrig *et al*, 2008) were grown in a 1:1 mixture of MEM and F-12 (both from Sigma), enriched with 10% fetal bovine serum (PanBiotech), 1% (Thermo Fisher Scientific, and 1% L-glutamine (PanBiotech). Cultures were maintained at 37 °C in a humidified incubator with 5% CO₂. The cells were transfected with either an empty pCEP4 vector (Invitrogen, negative control) or one of three C99-expressing constructs: pCEP4-spA4ct-DA-WT, -I45F, or -V50F. Transfections were carried out using Lipofectamine and Plus Reagent (Invitrogen) following the manufacturer’s instructions. Hygromycin B (300µg/ml; Thermo Fisher Scientific) was used for selection of stably transfected clones. The expression of C99 was confirmed in isolated clones, and those showing comparable expression levels were selected for downstream analysis.

### Collection of conditioned media

To collect secreted Aβ_1-42_, 5 ml of culture medium was added to 10cm dishes containing SH-SY5Y cells at ∼70% confluence. After incubation for 16 to 48 h, the conditioned media were harvested and cleared by centrifugation at 13,000 rpm for 1min at 4 °C. These conditioned media were either used directly or stored at –80 °C for additional experiments.

### Quantification of Aβ in conditioned media by ELISA

Soluble Aβ species released into the conditioned media were quantified via an enzyme-linked immunosorbent assay (ELISA) specific for human Aβ_42_ (Invitrogen). The samples and standards were prepared according to the manufacturer’s instructions. 96-well plates precoated with Aβ-specific capture antibodies were incubated with samples and calibrators, followed by detection with biotinylated secondary antibodies and a streptavidin-HRP conjugate. The colorimetric reaction was developed using TMB substrate and stopped with sulfuric acid. The absorbance was measured at 450 nm using a microplate reader. Aβ concentrations were calculated by interpolation from a standard curve generated via the synthetic Aβ peptides included in each kit. All samples were analyzed in technical duplicates, and the mean values are reported. The data are expressed as nanograms (ng) of Aβ per milliliter (ml) of CM.

### Cell viability and apoptosis assay

To evaluate the effects of Aβ on the cell viability of ENS cells, a CellTiter-Glo® Luminescent Cell Viability Assay (Promega) was performed according to the manufacturer’s instructions. In brief, dissociated ENS cells (40,000 per well) were plated in white, extracellular Matrigel (ECM, Sigma Aldrich)-coated (1:100), 96-well plates (Greiner) and cultured for 3 days under differentiation conditions. The cells were then treated with Aβ at 10nM, 100nM, 1μM, 2.5μM, or 10μM for 72h. CellTiter-Glo substrate (1:1) was added prior to the assay. Luminescence, proportional to adenosine triphosphate (ATP) levels as an indicator for metabolically active cells, was measured via a Tecan^TM^GENios® Microplate Reader. Additionally, a combined triple-staining approach was used to assess both cell viability and apoptotic status. Following treatment, the cells were trypsinized and washed with ice-cold PBS. For viability assessment, the cells were incubated with 2μM Calcein AM (Thermo Fisher Scientific) at 37 °C for 15min. After further washing, the cells were resuspended in binding buffer and stained for apoptosis using the eBioscience™ Annexin V Apoptosis Detection Kit PE (Thermo Fisher Scientific) following the manufacturer’s instructions. This included incubation with 5μL of Annexin V-PE for 12min at room temperature, followed by 7-Aminoactinomycin D (7-AAD staining. Flow cytometry was conducted within 4h using a BD FACS Canto II system. Populations of viable, early apoptotic, and late apoptotic cells were quantified.

### Neurite outgrowth

Neurite dynamics were monitored via the IncuCyte^®^ Live-Cell Analysis System 3 (Sartorius) to analyze neurite dynamics. Dissociated ENS cells (40,000 per well) were seeded into black clear-bottom 96-well plates (Greiner µClear®) and were immediately exposed to Aβ (10nM, 100nM, 1µM, 2.5µM, 10µM). The plates were scanned every 4 hours over a 72-h period using a 10x objective, and four images were acquired per well. NeuroTrack™ software was used to quantify neurite outgrowth on the basis of specific morphological parameters as described below: Cell-Body Cluster Parameters = Segmentation Mode (Brightness), Segmentation Adjustment (0.4); Cleanup = Hole Fill (0µm^2^), Adjust Size (0 pixels), min Cell Width (10µM); Cell-Body Cluster Filters = Area (min = 50µM^2^); Neurite Parameters = Filtering (None), Neurite Sensitivity (0.75), Neurite Width (1µM). The growth rate of neurites in each well was obtained by measuring the surface area covered by the neurites and is expressed as mm/mm^2^. Neurite growth was normalized to the initial time points (0h) and controls.

### Immunocytochemistry

For immunocytochemistry, dissociated ENS cells were fixed with 4% formaldehyde for 10min, permeabilized with 0.1% Triton X-100 for 10min, and blocked with 10% normal donkey serum in 0.01M PBS. The samples were incubated with primary antibody for 1h at room temperature (RT) in 0.01M PBS. To display the neurons, the cells were stained with class III beta tubulin (Tuj1, Biolegend, 1:500), and the synapses were visualized with vesicle-associated membrane protein 2 (Vamp2, Synaptic Systems, 1:500). The samples were washed three times in PBS and then incubated with the corresponding secondary antibodies (donkey anti-rabbit 594 and donkey anti-mouse 488, both Thermo Fisher Scientific, each 1:500 in 0.01M PBS) for 1h at RT. After further washing steps in 0.01M PBS (3x), the samples were washed once in distilled water. The cells were mounted on slides via fluorescent mounting medium with DAPI (Thermo Fisher Scientific). Imaging was performed using a Zeiss CellObserver Z1 and Axiovision software. To determine the percentage expression of the markers per image section, 10 images per condition were analyzed. The ratio of Vamp2 to Tuj1 signals was calculated to assess the synaptic density per neuronal area.

### MEA measurements

To measure the electrophysiological effects of Aβ on the neuronal activity of ENS cells, the MEAs 60MEA200/30iR-Ti from Multichannel Systems (MCS, Reutlingen) were used. Each array contains 60 electrodes with an interelectrode spacing of 200 µm and an electrode diameter of 30µm, along with an integrated reference electrode. The electrical activity of the neuronal networks was measured as extracellular field potentials. For the experiment, ganglia of ENS cells were isolated, allowed to proliferate for one day and then seeded at a density of 200,000 cells per chip due to the large surface area of the chip, which was precoated with a combination of poly-D-lysine (Merck) / ECM (1:100, Sigma Aldrich). After 3 days of differentiation, electrical activity was recorded using a MEA2100-Mini-System (MCS), which included a signal collector unit and interference board. During the recording, the headstage was kept in an incubator (37°C, 5% CO_2_). Electrical activity was acquired with MultiChannel Experimenter software (V. 2.21.1). Recordings were digitized at 10 kHz and filtered with a high pass Butterworth filter at a cutoff of 300 Hz and a low pass of 3,000 Hz to eliminate background activity and noise, which predominantly affect high and low frequencies, respectively. Recordings were processed offline with MultiChannel Analyzer software (V. 2.21.1.0). For signal detection, the threshold level was set with an average noise of five times the signal standard deviation on a per-channel basis (dead time of 3ms, pretrigger of 1.1 ms, and posttrigger of 2ms = 32 samples). Processed files containing whole data streams and detected signals (i.e., spike cutouts) were converted to HDF5 files in the MultiChannel DataManager (V. 1.14.10.23023) and read into a self-written MATLAB script (R2024b) using the McsMATLABDataTools Release 1.3.1. For spike analysis, the spike number per minute was calculated from spike cutouts. Following baseline measurements of spontaneous activity, Aβ peptides were applied at concentrations of 10nM, 100nM, and 1µM. Only lower-range concentrations were included in the analysis, as physiological levels of Aβ in the body typically fall within the nano-to picomolar range (Van Gool *et al*, 1995). The focus of this study was to investigate responses within the physiologically relevant range.

### Measurement of changes in intracellular Ca^2+^ levels

To evaluate intracellular Ca^2+^ levels, 40,000 ENS cells were seeded on ECM-coated (1:100) glass coverslips and differentiated for 3-5 days. The cells were loaded with 5µM Fluo-4 AM (Invitrogen) in the dark for 40min. Following the washing steps, the coverslip with the cells was fixed and aligned in the perfusion chamber RC-20 (Warner Instruments) for live-cell imaging. Measurements were performed under continuous perfusion with physiological imaging buffer (148mM NaCl, 5mM KCl, 2mM CaCl_2_, 1mM MgCl_2_, 10mM glucose, and 10mM HEPES [pH 7.38, all Thermo Fisher Scientific]), either alone or supplemented with Aβ (10nM, 100nM, or 1µM). Calcium imaging was conducted using a Zeiss CellObserver Z1 and the Axiovision software (Rel. 4.6.2.). Neuronal baseline activity was recorded for 600 seconds, followed by Aβ stimulation for an additional 600 seconds. To confirm neuronal responsiveness, 75 mM KCl was added as a positive control. Regions of interest were defined around fluorescent neuronal cell bodies. Calcium responses were quantified as F/F₀ (F: fluorescence at a given time; F₀: baseline fluorescence at time point 0), with calcium influx (ΔCa²⁺) calculated as (F − F₀)/F₀. To identify differences in calcium transients, the area under the curve (AUC) of the fluorescence intensity curve was calculated via custom-made Python script (V. 3.9). Four independent experiments were performed for each Aβ concentration. For each animal, the mean value of ΔCa^2+^ was calculated from all analyzed cells on a given coverslip.

### Motility assessment

To investigate the effect of Aβ on intestinal motility, a combined approach of mesenteric and luminal perfusion was used, as previously described (Schreiber *et al*, 2014; Gries *et al*, 2021). In brief, a 2.5cm segment of the jejunum was excised and mounted on Luer locks in an organ bath containing Tyrode buffer (130mM NaCl, 24.2mM NaHCO_3_, 11mM glucose, 4.5mM KCl, 2.2mM CaCl_2_, 1.2mM NaH_2_PO_4_, and 0.6mM MgCl_2_). Luminal perfusion was performed using a peristaltic pump at 0.2 ml/min, with the outflow tubing adjusted to a height of 3 cm to control luminal pressure. Simultaneously, the mesenteric artery was cannulated using a glass capillary, and Aβ was administered via a syringe pump (0.2 ml/min). In additional experiments, supernatants of transgenic Aβ-producing neuroblastoma cell lines were used to investigate the trigger for increased GI activity. Here, the gut was additionally perfused with 100nM MLA (Merck), a blocker of nicotinic acetylcholine receptors. The organ bath was continuously gassed with carbogen to maintain a physiological pH of 7.4 and held at 37°C using a heat exchanger. Preparations were equilibrated for 10min, followed by 10min of spontaneous movement recording and 10min of Aβ administration (100nM). Intestinal motility was recorded at 25 frames per second via video cameras. The data were analyzed using MotMap (www.smoothmap.org) and a custom LabVIEW script (LabVIEW 2019, National Instruments), as described previously (Schreiber *et al*, 2014; Gries *et al*, 2021).

### Ethics approval and consent to participate

All procedures involving animal tissue were performed in accordance with institutional and national guidelines. The intestinal tissue was obtained from euthanized WT C57BL/6J mice in accordance with the regulations of the Landesuntersuchungsamt of Rhineland-Palatinate (Koblenz). No animal experiments were performed as part of this study. The removal of organs was approved within the institutional framework for the use of animal by-products for scientific purposes.

### Statistical analyses

Statistical analyses were performed using GraphPad Prism 8. Data were normalized to the untreated control (viability, apoptosis, immunocytochemistry, motility to baseline activity, and neurite outgrowth additionally to time point 0), whereas MEA and calcium imaging data are shown as raw values. The normality of the data distributions was assessed via the Shapiro‒Wilk test. For normally distributed data, differences between groups were evaluated using the Student’s t test. For normalized datasets, comparisons were made using one-way ANOVA or a one-sample t test, with the latter testing against the control condition with a hypothetical mean of 100. The results are expressed as the means ± standard deviations (SDs). Statistical significance was set at *p* ≤ 0.05 (*), *p* ≤ 0.01 (**), and *p* ≤ 0.001 (***).

## Acknowledgements

This work was partly funded by the “*Offene Digitalisierungsallianz Pfalz*” project, which is part of the federal-state initiative “Innovative University.” Furthermore, funding was provided by the EU Joint Programme – Neuro-degenerative Disease Research (JPND) and BMBF grant EURO-FINGERS 01ED2003.

## Author’s contributions

M.G., H.P. and K-H.S. conceived the project, designed and supervised all the experiments, analyzed and interpreted the results and wrote the manuscript. S.S helped generate and analyze all MEA and Ca imaging data, wrote the MatLab and Python scripts, and helped with sample preparation. A.C. performed and assisted with all motility studies. S.R. assisted with sample preparation and Ca-Imaging experiments. M.B. helped analyze the data and collect the statistics. M.M. helped with the calcium imaging experiments. M.O.W.G. and T.H. contributed to the additional perfusion experiments with SH-SY5Y supernatants. All the authors discussed the results and commented on the manuscript.

## Availability of materials

All datasets and raw imaging files generated during this study are available from the corresponding author upon reasonable request.

